# Surface potential as a strong early osteogenic trigger via mechanotransduction and calcium accumulation

**DOI:** 10.64898/2026.04.26.720950

**Authors:** Sara Martin-Iglesias, Yaiza R. Varela, Paula Rodriguez-Lejarraga, Lucia Jiménez-Rojo, Cristina Eguizabal, Noemi Jiménez-Rojo, Juan Anguita, Ana M. Aransay, Senentxu Lanceros-Méndez, Unai Silvan

## Abstract

Analyzing the differentiation potential of cells in contact with newly developed materials is essential for assessing their ability to integrate into biological tissues and promote functional regeneration. Material properties such as rigidity, topography, and wettability significantly influence stem cell differentiation and are therefore optimized in implants. In this context, surface potential has been repeatedly, albeit inadvertently, shown to enhance osteogenesis. Here, we demonstrate that this surface property modulates cellular mechanosensing by altering the cell’s perception of substrate rigidity. Specifically, we show that human bone marrow-derived mesenchymal stem cells (hBM-MSCs) on surfaces with a net zero charge, coated with collagen type I, exhibit characteristics typical of cells adhering to compliant substrates. Conversely, mesenchymal stem cells on polarized surfaces activate mechanoresponsive pathways that promote osteogenesis, as evidenced by large spreading areas, enhanced contractility, and Yes-associated protein (YAP) translocation into the nucleus. Furthermore, our data suggest that negative net surface potentials lead to the local accumulation of calcium ions, which further facilitates osteogenic differentiation. Collectively, our findings reveal that biomaterials’ surface potential, a previously uncharacterized mediator of cellular mechanotransduction, should be considered in the design of next-generation biomaterials for tissue regeneration applications.

## Introduction

The design of new biomaterials for tissue engineering requires a precise understanding of their interactions with biological tissues so that characteristics supporting specific cellular behaviors can be maximized. For instance, in vitro analysis of the impact of nano- and micro-topography on osteogenesis has enabled the development of bone implants with optimal roughness and improved osseointegration (1–3). Similarly, other surface properties, such as wettability (4), ligand type and density (5–9), and deformability (10), have been identified as potent regulators of cell behavior and thus optimized for specific applications. Overall, the transduction of these environmental signals relies on focal adhesions (FAs) that connect the extracellular space with the cell’s contractile actomyosin machinery (11, 12). The binding of integrins in FAs to the extracellular matrix (ECM) depends on ligand properties and ultimately determines intracellular tension, which propagates to the nucleus and activates signaling pathways (13–15). Central to this response is the Yes-associated protein (YAP or YAP1), a transcriptional co-regulator within the Hippo signaling pathway. Upon mechanical activation, YAP translocates to the nucleus and binds to enhancer elements, promoting the expression of target genes, including regulators of the cell cycle, migration, and differentiation (16, 17).

Over the last decade the impact of surface potential on stem cell lineage commitment has been analyzed using polymeric (18–20), ceramic (21–23) and metallic materials (24–26). Research has consistently shown that polarized surfaces are more effective at promoting osteogenesis than non-polarized ones; however, the polarity that maximizes osteogenic differentiation has varied depending on the study. It has been shown that the accessibility of cell adhesion motifs and the subsequent cell response strongly depend on the physicochemical properties of the surface where collagen is deposited (4, 27). Similarly, analysis of fibronectin and collagen deposited on poled (with either positive or negative surface charge) and non-poled (with net zero surface charge) polyvinylidene difluoride (PVDF) surfaces suggests that surface potential induces conformational changes that promote the exposure of cryptic cell binding sites (20, 28). Furthermore, since the distribution of charged amino acids on the surface of the protein is not uniform, proteins can physically organize themselves so that regions with opposite charges bind to the substrate, leaving regions with the same charge sign as the original surface exposed to the culture medium. Nevertheless, it is still unclear how these surface-driven structural changes impact cell behavior, or what cellular mechanisms actually drive surface potential-induced differentiation.

Herein, we systematically analyzed the response of human bone marrow-derived mesenchymal stem cells (hBM-MSCs) to PVDF substrates with different permanent surface electric charges but equivalent roughness, ligand densities, wettability, and rigidity. Our results reveal differences in the apparent stiffness sensed by hBM-MSCs, depending on the net surface charge of the substrate to which they adhere. Specifically, on neutral (non-poled) surfaces, hBM-MSCs show morphological features and mechanoresponsive signaling characteristic of cells adhering to compliant surfaces. Conversely, on positively and, especially, negatively polarized surfaces, cells activate rigidity-mediated pathways, thus displaying a higher tendency toward osteogenic differentiation (10). Moreover, our data suggest that negatively charged surfaces promote the local accumulation of calcium ions, further enhancing osteogenic differentiation at longer culture times. This study is the first to describe the biological mechanisms governing the interaction of cells with surfaces of varying surface charges, and it highlights the importance of considering this physical characteristic in the development of improved biomaterials for tissue regeneration applications.

## Results

Cells respond to various material properties, including topography, rigidity, hydrophobicity and ligand distribution (29, 30). To exclude the effect of incidental properties of the materials, in our experiments we used commercial polyvinylidene difluoride (PVDF) films with varying surface electric charges, but identical chemical composition and wettability (Figure 1a-c). The different surface electric potentials are achieved through a polarization process of the films, which results in polar β-phase PVDF (β-PVDF) with either a permanent net positive (PVDF (+)), negative (PVDF (−)) or neutral average surface charge (PVDF (non-poled)) (31). To avoid any bias caused by different ligand densities, we ensured that collagen type I and fibronectin coatings were equivalent across all conditions (Figure 1d).

**Figure 1.**
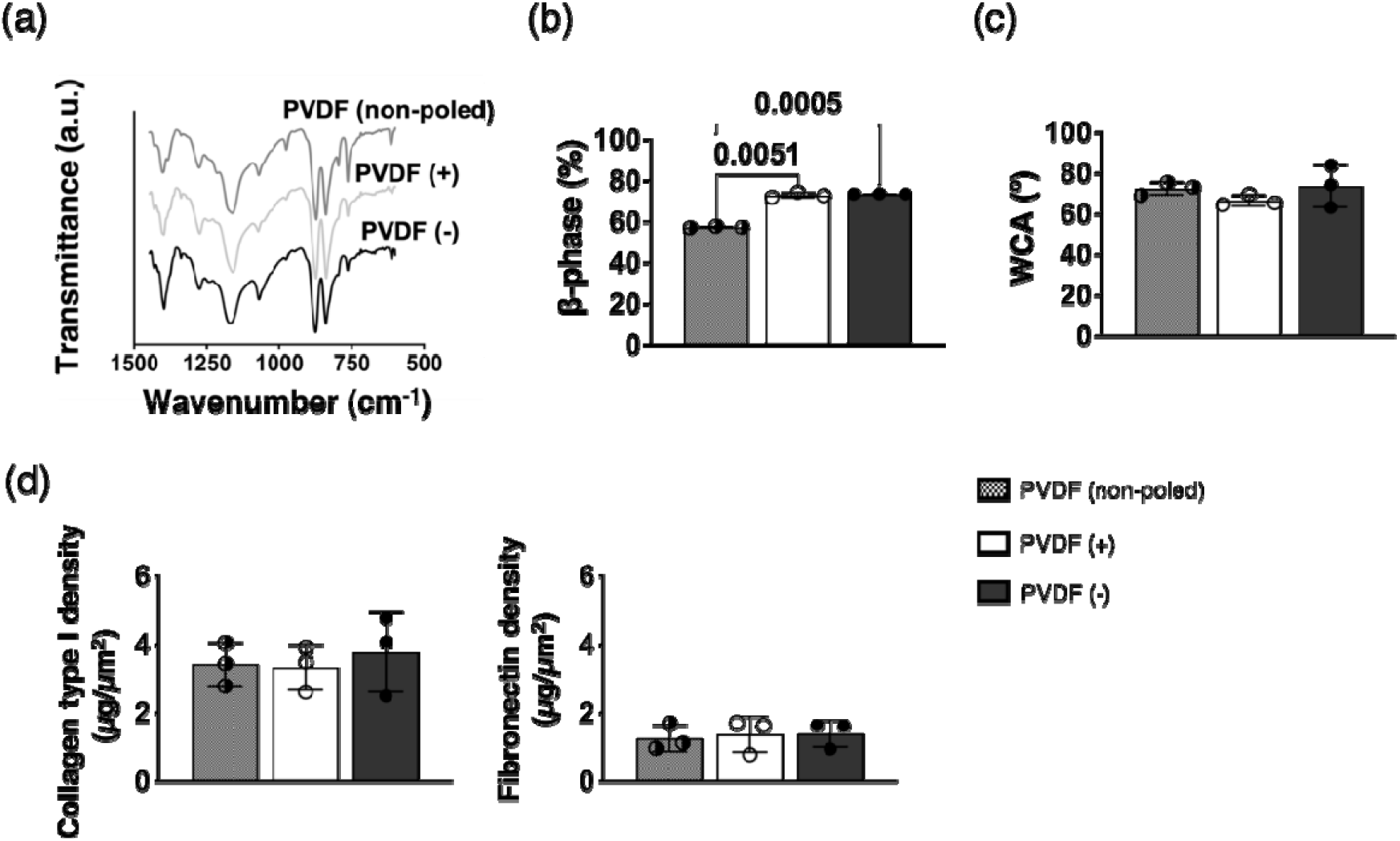
Material characterization. (a) FTIR spectra of poled and non-poled β-PVDF films and (b) calculated β-phase content. (c) Water contact angle (WCA) of the bare surfaces. (d) Collagen type I (left panel) and fibronectin (right panel) density on the different β-PVDF substrates. Results are expressed as mean ± SD (n=3). Only p-values <0.1 are shown.

First, we evaluated how effectively each surface supports osteogenesis. An experimental condition with silica glass coverslips was included, as these surfaces are often used as positive controls in osteogenic differentiation assays (32). After seven days of osteogenic induction we measured the highest alkaline phosphatase (ALP) activity in hBM-MSCs adhering to β-PVDF with negative electric potential coated with collagen type I, being the control condition (glass coverslips) equally osteogenic (Figure 2a). In turn, fibronectin coating rendered glass coverslips and all β-PVDF surfaces equally osteogenic, regardless of their net surface charge (Figure 2b).

**Figure 2.**
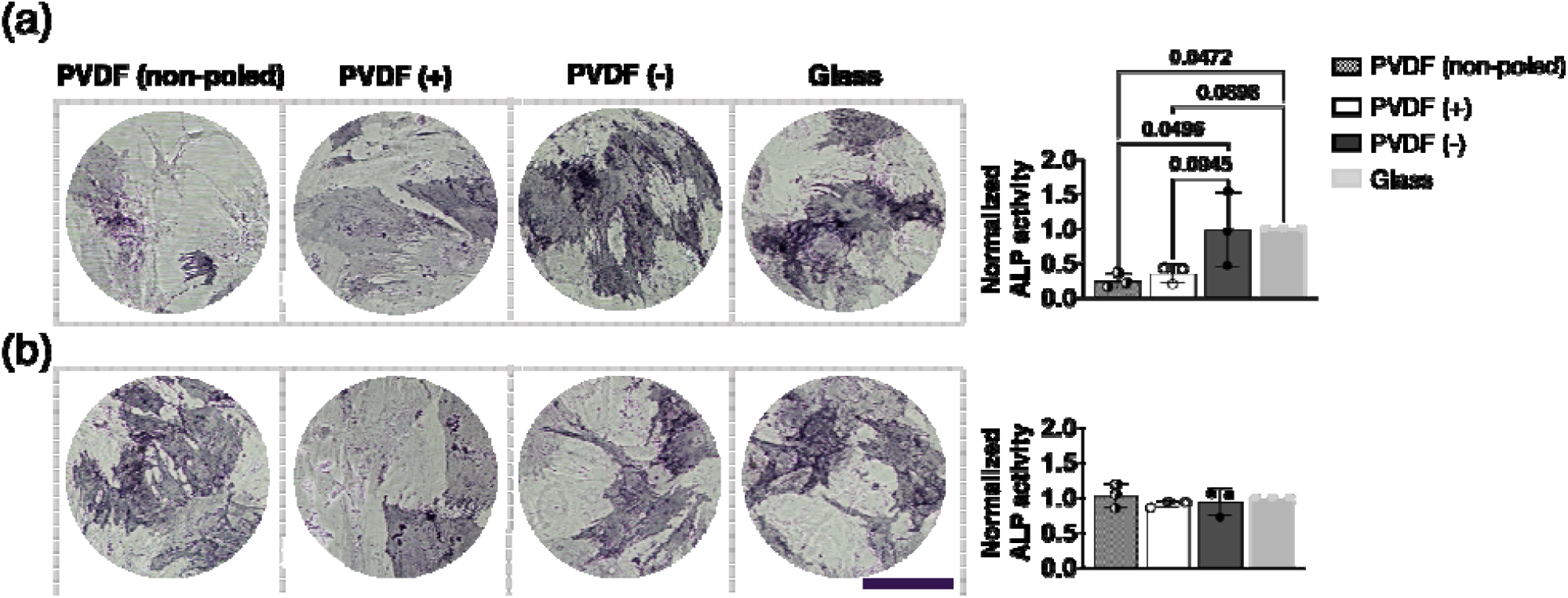
Alkaline phosphatase activity. Colorimetric staining and quantification of ALP activity in hBM-MSCs cultured for seven days on collagen type I-coated (a) and fibronectin-coated (b) substrates. Results are expressed as mean ± SD (n=3). Data were analyzed assuming a Gaussian (normal) distribution and equal variances across groups; statistical differences were assessed using one-way ANOVA. Only p-values <0.1 are shown. Scale bar represents 200 µm.

Since cell shape and fate are closely related (33), we analyzed the morphological features of hBM-MSCs adhering to the substrates by means of immunofluorescence. After 24 hours and four days culture on the β-PVDF films, hBM-MSCs display morphologies largely dependent on the electric surface charge and coating used. Specifically, the footprint and nuclear projected area of cells on collagen type I–coated negatively and positively poled surfaces were larger than on non-poled ones (Figure 3a and supplementary figure 2). Simultaneously, the FA count per cell and FA density appeared upregulated. Meanwhile, on fibronectin-coated surfaces, cells did not exhibit morphological changes dependent on the surface charge (Figure 3b and supplementary figure 3).

**Figure 3.**
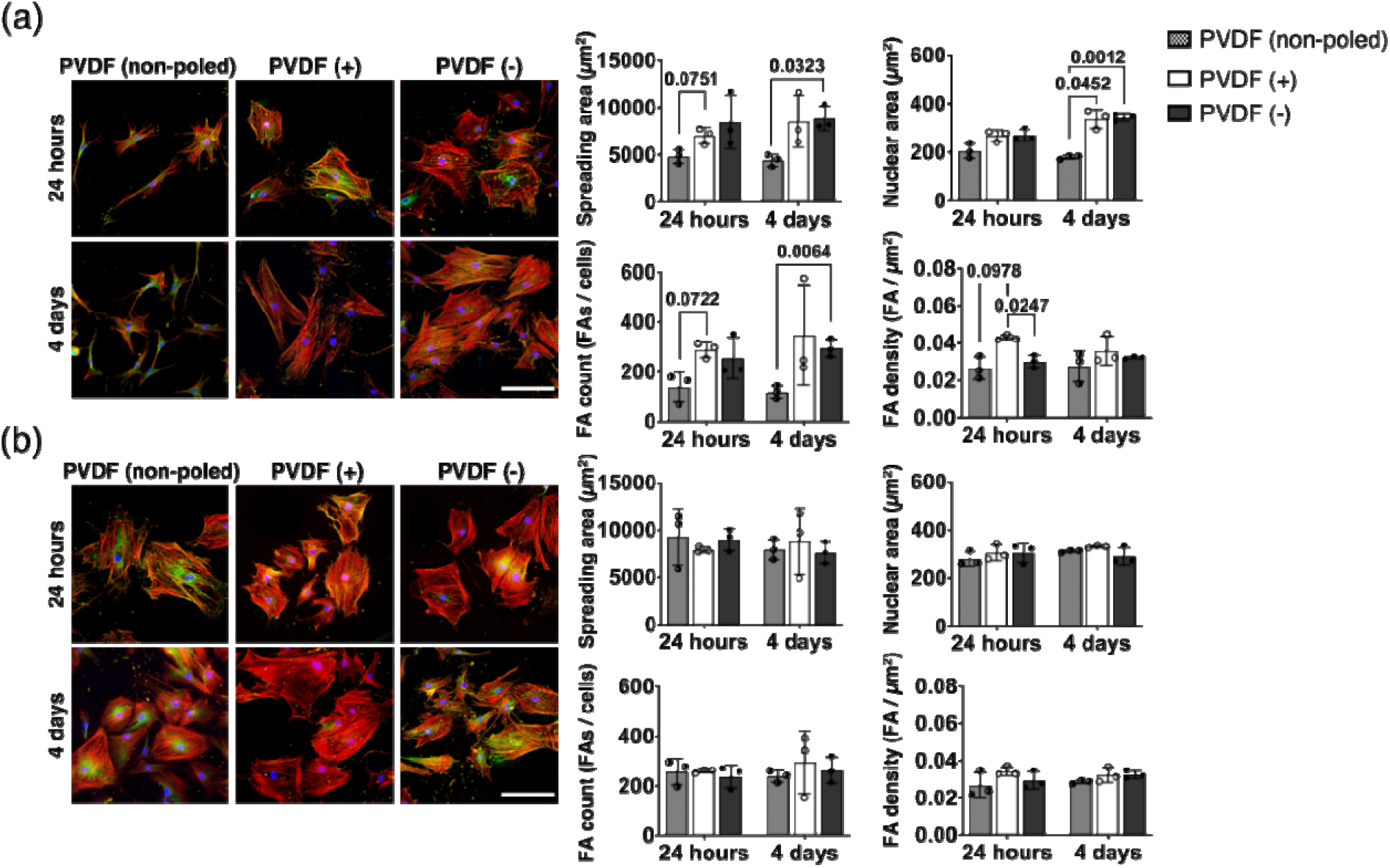
Morphological analysis of hBM-MSCs cultured on β-PVDF films of varying surface potential. Fluorescence images of hBM-MSCs cultured on β-PVDF films coated with either collagen type I (a) or fibronectin (b) and corresponding morphological features. Nuclei are stained in blue, vinculin in green and F-actin in red. Results are expressed as mean ± SD (n=3). Only p-values <0.1 are shown. Scale bars represent 200 µm.

Rho-associated protein kinases I and II (ROCK I and II) are critical regulators of cytoskeletal dynamics via their interaction with downstream effectors, such as Myosin phosphatase target subunit 1 (MYPT1), which inhibits myosin light chain (MLC) phosphatase. Through MLC phosphorylation, modulation of F-actin organization, and FA turnover, the ROCK-MYPT1 axis impacts cell shape and differentiation, particularly in MSCs (34, 35). Thus, we quantified the relative MYPT1 phosphorylation in hBM-MSCs adhering to the different surfaces and found higher values in hBM-MSCs adhering to positive and, especially, negative β-PVDF substrates coated with collagen type I (Figure 4a). In contrast, the phosphorylation levels of MYPT1 did not show any differences on fibronectin-coated films, regardless of the net surface charge of the substrates (Figure 4b).

**Figure 4.**
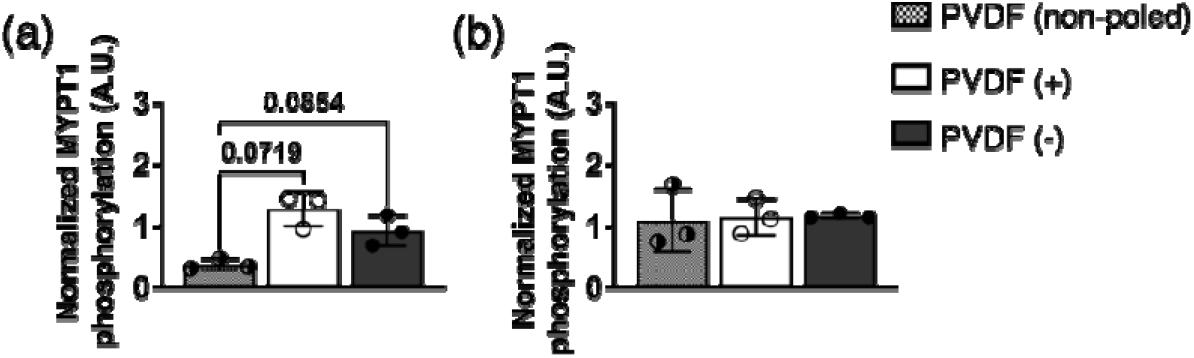
Signal transduction. MYPT1 phosphorylation in hBM-MSCs cultured on collagen type I-coated (a) and fibronectin-coated (b) β-PVDF films of varying surface potential. Results are expressed as mean ± SD (n=3). Only p-values <0.1 are shown.

Intracellular tension and contractility have been shown to promote the translocation of YAP into the nuclear compartment, which in turn drives cell fate determination (36). In our experimental setup, nuclear translocation of YAP was highest in hBM-MSCs adhering to collagen-coated β-PVDF surfaces with either negative or positive net charge, compared with those adhering to non-poled surfaces (Figure 5a). In contrast, on fibronectin-coated substrates, YAP was preferentially detected in the nucleus, regardless of the surface potential (Figure 5b).

**Figure 5.**
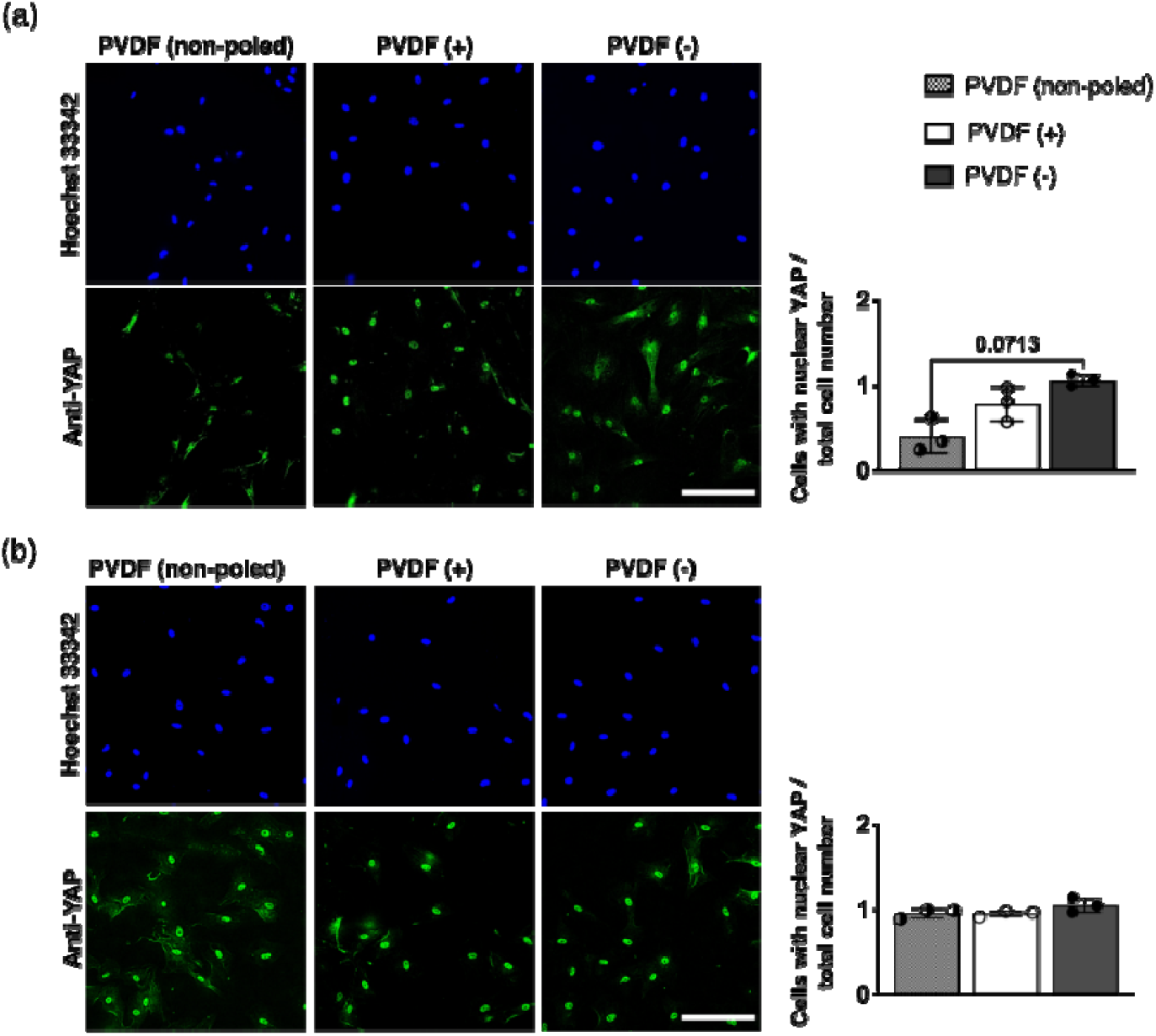
Nuclear translocation of YAP. Fluorescence images of hBM-MSCs cultured on β-PVDF films coated with either collagen type I (a) or fibronectin (b) immunostained against YAP and counterstained with Hoechst 33342, and the corresponding quantifications (right panels). Scale bars represent 200 µm. Results are expressed as mean ± SD (n=3). Only p-values <0.1 are shown.

In the nucleus, YAP interacts with TEAD family transcription factors and promotes the expression of specific genes. Thus, we performed RNA sequencing of hBM-MSCs cultured for 24 hours and four days on the substrates coated with collagen type I. The comparison of the gene expression patterns of hBM-MSCs cultured on non-poled β-PVDF and on positively and negatively charged β-PVDF for 24 hours revealed 598 and 407 differentially expressed genes, respectively. Of these, 265 genes were uniquely regulated in cells on the positive surfaces and 74 on the negative surfaces (Figure 6a). Following four days of culture, the number of differentially expressed genes decreased to 320 for hBM-MSCs adhering to non-poled compared with positive β-PVDF and to 352 compared with negative. In this case, the number of genes uniquely regulated in cells on positively charged surfaces was 115, while it increased to 147 genes on negatively charged surfaces (Figure 6b). Notably, hBM-MSCs cultured on positively and negatively charged surfaces at either time point were transcriptionally similar in terms of their overall gene expression profiles (Supplementary figure 4a and b) and no genes were differentially expressed between them (Supplementary figure 4c). Gene Set Enrichment Analysis (GSEA) revealed a significant suppression of gene sets related to cytoskeletal organization and morphogenesis in hBM-MSCs cultured on β-PVDF (non-poled) compared with those cells on surfaces with net charge (Figure 6c and d).

**Figure 6.**
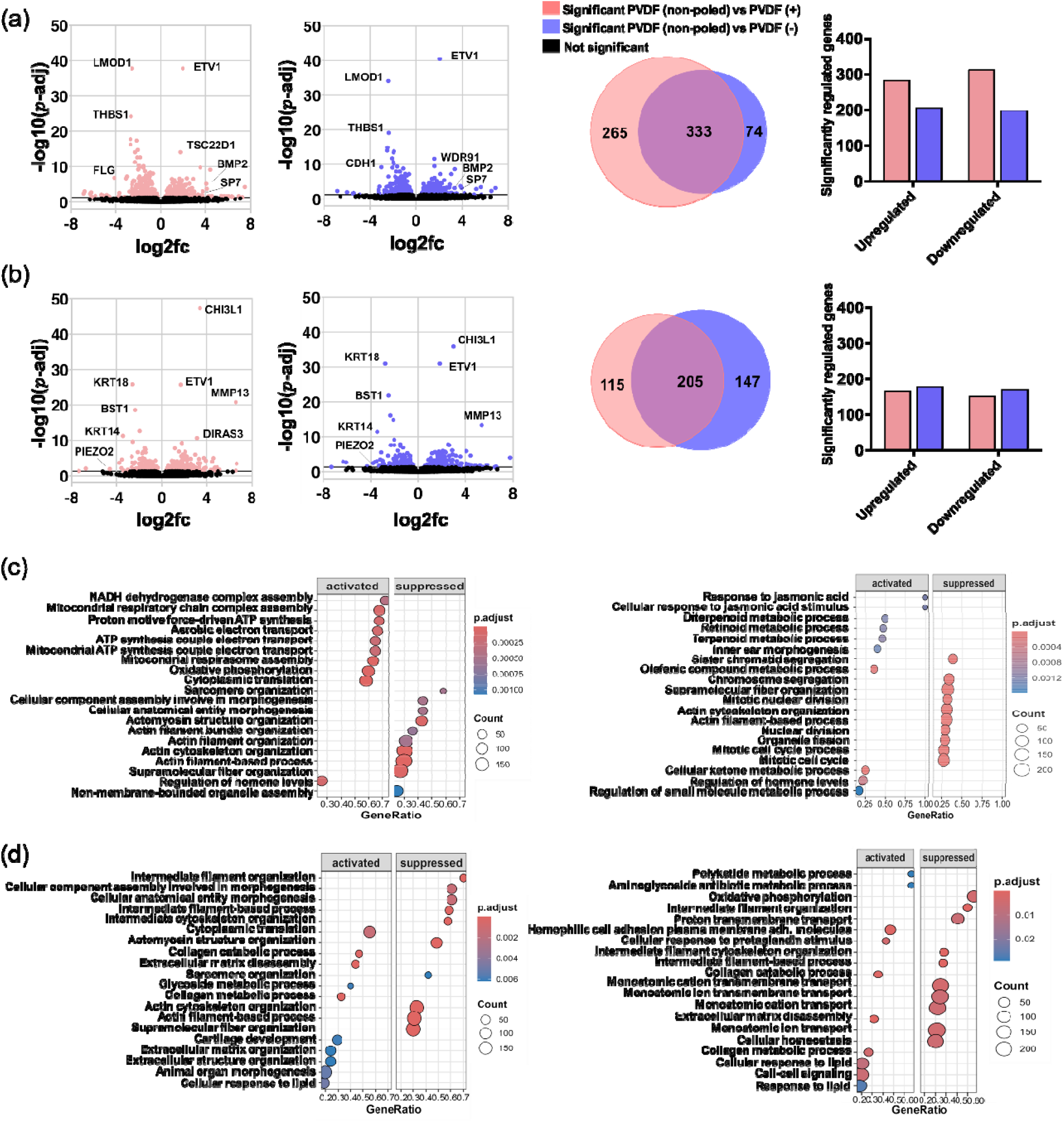
RNA sequencing. Volcano plots with the −log 10 (p-value) plotted against their respective log 2 (fold change) of genes differentially expressed in hBM-MSCs, Venn diagrams and histogram plots showing genes that are down- or upregulated (p<0.05) in hBM-MSCs cultured on the indicated surfaces for 24 hours (a) or four days (b). Circle area in a and b is proportional to the number of genes. GSEA of the cells cultured on the different surfaces for 24 hours (c) and 4 days (d).

The results suggest that cells sense varying rigidities depending on the net surface charge of the substrates; consequently, we performed experiments in which, after collagen-coating, the protein interface was chemically stiffened using glutaraldehyde crosslinking (37). Under these conditions, the morphological features of hBM-MSCs adhering to the β-PVDF films, including footprint, adhesion sites per cell, and nuclear area, remained comparable across all polarities at the 24-hour time point (Figure 7a and b, and supplementary figure 5). However, after four days in culture, small differences were observed between polarities though these variations were less pronounced than in the experiments using native collagen type I. Similarly, the previously observed differences in MYPT1 phosphorylation driven by surface charge were absent in the surfaces with fixed collagen (Figure 7c), and a similar relative number of cells with nuclear YAP were quantified in β-PVDF substrates, regardless of their polarity (Figure 7d). Furthermore, treatment of hBM-MSCs with the cell-permeable ROCK inhibitor Y-27632 largely reduced the morphological differences between cells cultured on different β-PVDF substrates (Figure 8a and b, and supplementary figure 6) and the ratio of cells on poled surfaces exhibiting nuclear YAP (Figure 8c and d).

**Figure 7.**
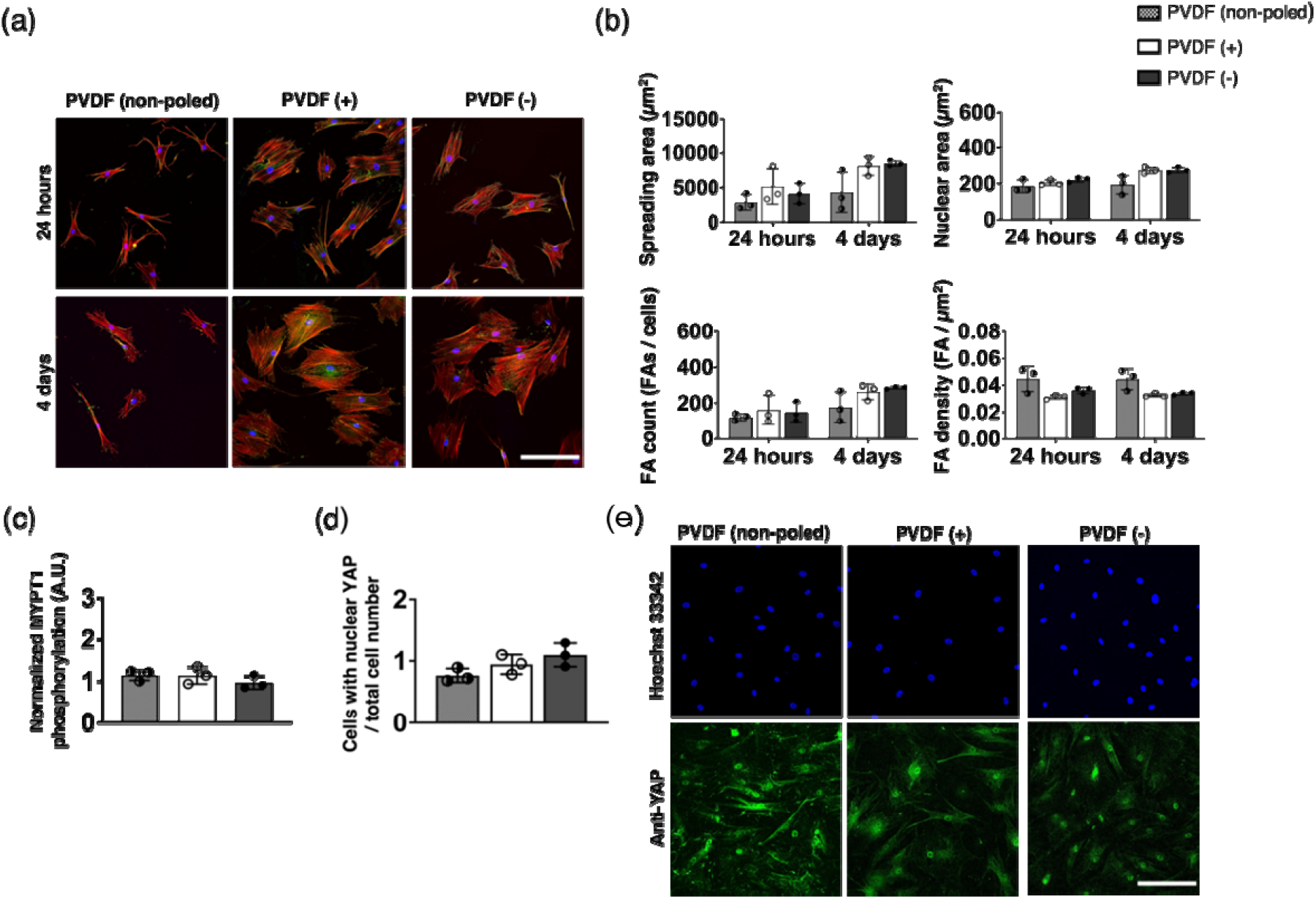
Stiffening of the collagen type I coating. (a) Immunostaining of hBM-MSCs cultured on the different β-PVDF surfaces with glutaraldehyde-crosslinked collagen type I coating and corresponding morphologic analysis (b). MYTP1 phosphorylation (c) and YAP translocation (d and e) in hBM-MSCs cultured on the substrates. Only p-values <0.1 are shown. Scale bars represent 200 µm. Results are expressed as mean ± SD (n=3).

**Figure 8.**
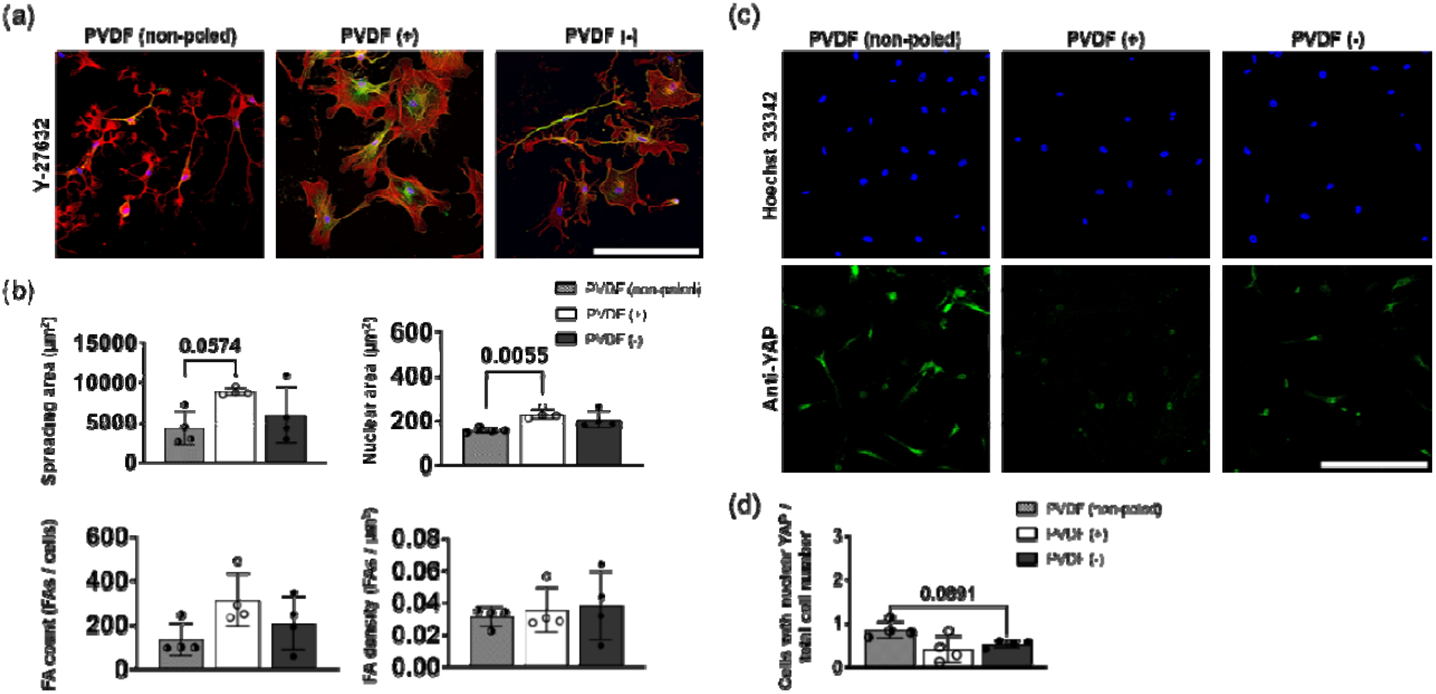
ROCK inhibition. Immunofluorescent images (a) and corresponding morphologic analysis (b) of hBM-MSCs treated with the ROCK inhibitor Y-27632 for 24 hours. YAP immunolocalization (c) and translocation (d) in treated cells. Only p-values <0.1 are shown. Scale bars represent 300 µm. Results are expressed as mean ± SD (n=4).

Previous studies have reported that undifferentiated hBM-MSCs exhibit asynchronous oscillations of intracellular calcium ions (Ca^2+^), which are not uniformly present throughout the population (38). These spontaneous spikes vary in frequency and pattern among individual cells and depend on the presence of extracellular calcium ions (39). To identify cells exhibiting such Ca^2+^ oscillations, we established a criterion of at least three transient increases in Fluo-4 fluorescence intensity greater than 75% above basal levels over the imaging period— approximately equivalent to 0.2 events/minute (Figure 9a and b). Using this threshold, we found that, regardless of substrate charge, MSCs showed comparable frequencies of Ca^2+^ transients across all culture conditions (Figure 9c). However, the mean duration of individual Ca^2+^ spikes, defined as the time the fluorescence signal remained above the intensity threshold, was significantly longer in hBM-MSCs adhering to β-PVDF (−) (Figure 9c). To determine the downstream effects of the observed calcium oscillations, we further analyzed the phosphorylation status of ERK1/2. Western blot analysis revealed increased ERK1/2 phosphorylation in hBM-MSCs on β-PVDF (−) compared with those on β-PVDF (+) or β-PVDF (non-poled) (Supplementary figure 7a and b).

**Figure 9.**
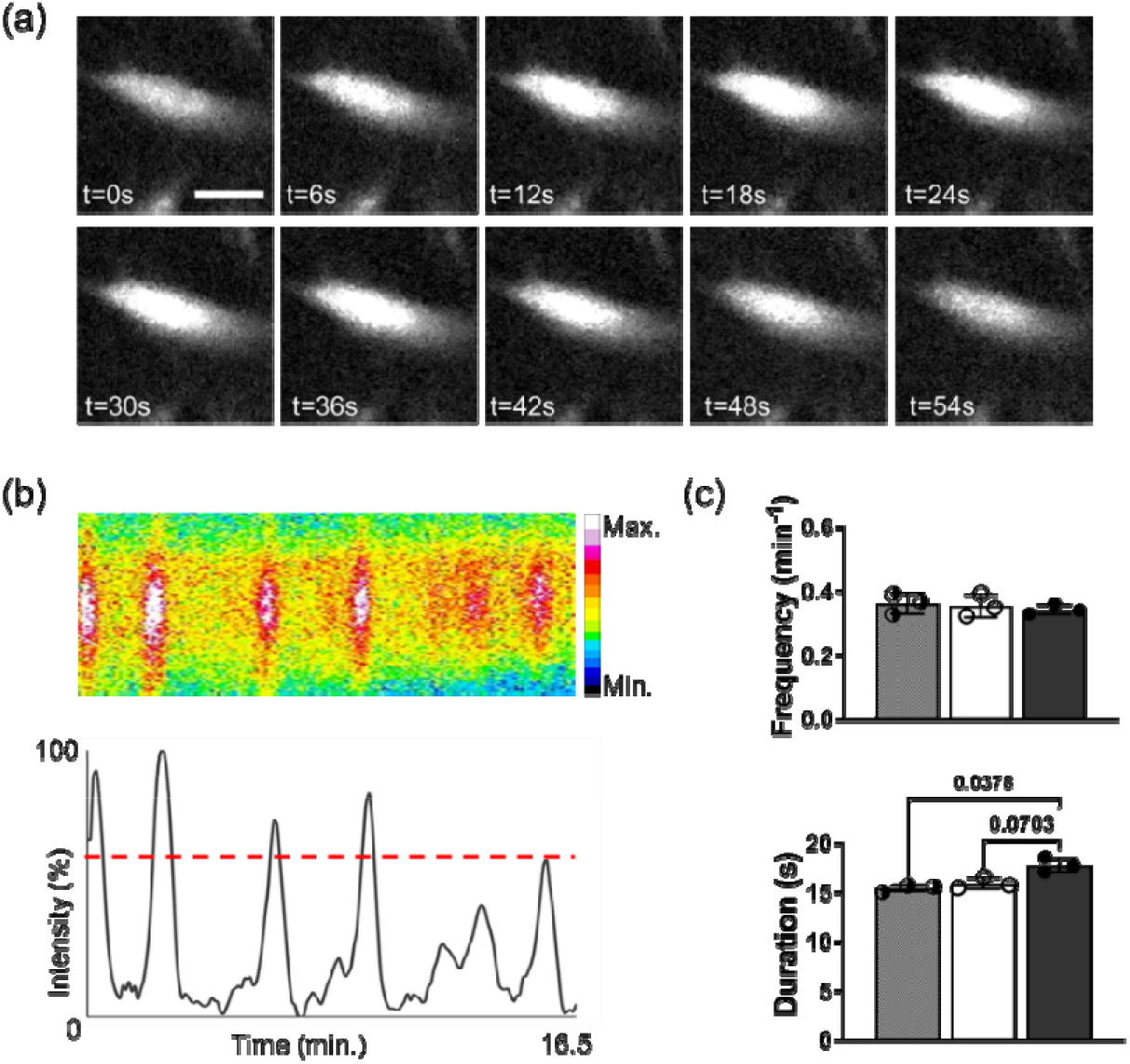
Calcium signaling. (a) Representative image series of a Ca^2+^ event in an MSC adhering to β-PVDF (−). (b) Kymograph (upper panel) and corresponding histogram (bottom panel) showing the intensity fluctuation of a hBM-MSC cultured on β-PVDF (−). (c) Peak frequency (upper plot) and mean duration of the events (bottom plot). Only p-values <0.1 are shown. Scale bar represents 40 µm. Results are expressed as mean ± SD (n=3).

## Discussion

New materials that can promote osteogenic differentiation are regularly reported; however, their descriptions generally lack a systematic assessment of the relative contributions of specific physicochemical properties to the observed biological response and, in most cases, fail to explain the underlying mechanisms. While such descriptive studies provide valuable data and facilitate the identification of promising candidates for specific purposes, determining critical variables and driving forces is necessary for the evidence-based design of materials with predefined biological activities (40). In fact, systematic in vitro analysis of the biological response to material properties such as topography (32, 41), rigidity (40) and cell adhesion (3), has enabled the design of biomaterials with enhanced abilities to support specific cellular behaviors. Similarly, several studies indicate that surface electric potential is a potent regulator of lineage commitment (24, 42, 43). Nevertheless, the underlying mechanisms and involved transduction signaling pathways remain largely undescribed, so this property is generally neglected.

In this study, we performed a comprehensive analysis of the response of hBM-MSCs to surfaces with different surface potentials but otherwise identical properties. To this end, we used PVDF, an electroactive material that displays excellent biocompatibility and can be produced with different permanent surface polarities (14). We observed that the early response of hBM-MSCs to electrically charged β-PVDF surfaces is similar to that observed on rigid substrates, characterized by large spreading areas, increased phosphorylation of MYPT1, and nuclear translocation of YAP (44). This response ultimately leads to increased osteogenic differentiation, as previously reported (18). In turn, cells on non-poled β-PVDF display features characteristic of cells adhering to compliant materials (10). These findings suggest a weaker interaction between collagen and the non-poled PVDF surface; this increased matrix mobility leads cells to perceive the microenvironment as mechanically softer. The suppression of polarity-driven morphological variations due to chemical fixation of the protein interface further support differences in collagen tethering as a driving mechanism (45). On the other hand, the insensibility to surface polarity observed in experiments using fibronectin coating may be attributed to the activation of specific integrin-mediated signaling (46–48) and higher protein-material binding forces (49). Interestingly at short culture times hBM-MSCs on positively and negatively-poled surfaces did not display significant differences in their gene expression profiles. This suggests that surface potential-driven osteogenic differentiation is dependent on further signaling mechanisms that require longer culture times to trigger lineage commitment. In this regard, besides physicochemical cues, osteogenic differentiation is largely determined by biochemical factors present in the cell microenvironment, including growth factors and inorganic ions (50). Among the latter, calcium is the prevalent cation in bone, constituting approximately 40% of the bone mineral by weight in the form of hydroxyapatite crystals (51). These calcium phosphate complexes provide structural rigidity while serving as mineral reservoirs (52). The balance between calcium deposition by osteoblasts and release by osteoclasts determines the local Ca^2+^ concentration in vivo, which plays a central role in the generation and modulation of intracellular calcium oscillations, both by influx from the extracellular environment and by release from internal stores such as the endoplasmic reticulum. In turn, the fluctuations mediate the activation of osteogenic differentiation pathways in osteoprogenitor cells (53). Similarly, moderate presence of Ca^2+^, at a slightly higher concentration than in normal DMEM medium (1.8 mM), promotes proliferation and maturation of osteoblasts in vitro, while higher concentrations (> 10 mM) are cytotoxic (54– 56). Extensive published literature documents the accumulation of counterions on poled surfaces (57, 58). Our data suggest that the accumulation of Ca^2+^ in the vicinity of the cells (59) contributes to longer intracellular calcium oscillations that upregulate ERK phosphorylation and signaling pathways related to hBM-MSC differentiation. This biochemical difference, together with the increased mechanical rigidity sensed by the cells adhering to polarized surfaces ultimately results in the higher osteogenic activity of hBM-MSCs on the negatively charged surfaces.

Silica glass coverslips and tissue culture plastic (TCP) are commonly used as controls for the in vitro estimation of the potential of biomaterials to support osteogenesis. Interestingly, MSCs cultured on these substrates often demonstrate equivalent or even superior osteogenic differentiation potential compared with those adhering to the biomaterial surfaces under investigation (32, 43, 60–65). Glass coverslips are primarily composed of silica (SiO_2_) and boric oxide (B_2_O_3_). When placed in an aqueous solution, the silanol groups on its surface dissociate, causing the surface to acquire a net negative electrical charge. Meanwhile, tissue culture vessels are generally produced in polystyrene and rendered hydrophilic using corona discharge to promote cell adhesion (66–68). This treatment also makes the surface negatively charged when immersed in aqueous solutions (69, 70). Our work provides an explanation for the superior pro-osteogenic properties of these surfaces by uncovering the dual effect of negative surface charge on MSC differentiation. Given that TCP and glass coverslips frequently outperform the osteogenic potential of materials specifically designed with optimized surface properties, including wettability, roughness and composition (32, 43, 60), it is likely that surface potential plays a dominant role in regulating lineage commitment by affecting MSC spreading, mechanotransduction and Ca^2+^ availability.

Further work in the field should focus on identifying the specific ion channels mediating lineage commitment driven by surface potential (71) and on systematically evaluating how variations in the magnitude and spatial distribution of the surface charge impact this response. Additionally, future studies should elucidate if the dynamic changes in surface potential, characteristic of piezoelectric materials already used in tissue engineering, impact cellular behavior though mechanisms similar to those identified in the current study, and how cells process these complex stimuli into a biological response. This knowledge will enable the rational design of biomaterials with properties tailored to specific tissue engineering applications.

## Conclusion

In the current study we demonstrate that surface electric potential acts as a key factor in modulating stem cell lineage commitment through a process mediated by the interaction between cell mechanosensitivity and intracellular calcium signaling. By adjusting its electrical state, PVDF can actively guide and stimulate specific cellular responses. Ultimately, these insights provide a new material property that should be considered for the design of next-generation materials for tissue engineering applications.

## Materials and methods

### Fourier-transform infrared spectroscopy (FTIR) and polar β-phase content estimation

To estimate the polar β-phase content in the different PVDF substrates, Fourier-transform infrared spectroscopy - attenuated total reflectance mode (FTIR-ATR) measurements were carried out using a Jasco FT/IR-6100 spectrometer (Jasco, Japan). Briefly, spectral data were acquired in the 600–4000 cm^−1^ range at a resolution of 4 cm^−1^, and each spectrum resulted from averaging 64 individual scans. Three separate measurements were taken per sample. Bands at 766 cm^−1^ and 840 cm^−1^ were used to calculate the relative of the β-phase fraction as previously reported (72).

### Water contact angle (WCA) estimation

Films were positioned horizontally and leveled on the goniometer (Osscila, Netherlands) stage to ensure accurate droplet shape and reproducible image acquisition. A 5□µL droplet of deionized water was deposited onto the film surface, and lateral images were captured within 5 seconds after droplet deposition. Measurements were performed on at least three regions of the sample. Contact angles were analyzed using the “Contact Angle” plugin in Fiji (ImageJ).

### Substrate preparation

Commercial non-poled (PVDF-MDO-100, PolyK Technologies), with an average surface potential of 0 V, and poled β-PVDF films (11027826-00, TE Connectivity), with average positive (6 V) or negative (−4 V) surface charge (73), were cut into 15 mm diameter discs, sterilized with 70% ethanol and washed with PBS (10010-031, Gibco) and sterile water for 5 minutes each. For protein coating, polarized surfaces were incubated for 1 hour in an inverted position over a drop containing either 50 µg/ml rat tail type I collagen (A10483-01, Gibco) or fibronectin from bovine plasma (1030-FN-01M, R&D Systems). In turn, for non-poled substrates, type I collagen and fibronectin concentrations were 570 µg/ml and 100 µg/ml, respectively. Subsequently, coated substrates were washed with PBS to remove unbound proteins.

To quantify the amount of protein deposited on the surfaces a Bicinchoninic acid assay (BCA; 71285-3, Sigma-Aldrich) was used following the manufacturer’s instructions. Briefly, substrates were incubated in 400 µL of BCA working solution with copper reagent for 30 minutes at 37°C and the optical density (OD) of the resulting colorimetric solution was measured at 562 nm in a plate reader (Tecan Infinite MNano+). Known protein concentrations were used to create a standard curve and calculate the amount of protein absorbed per mm^2^ of culture area.

For the matrix stiffening experiments (37) collagen-coated substrates were submerged in a 2% (w/v) glutaraldehyde solution (G5882, Sigma-Aldrich) for 15 minutes at room temperature. Samples were then washed 3 times with PBS for 5 minutes, and immediately used in the in vitro tests.

### Alkaline Phosphatase Assay

hBM-MSCs (Cytion, 300665) were seeded at an approximate density of 5.000 cells/cm^2^ on the different surfaces in basal medium (DMEM; 11995-065, Gibco), 10% fetal bovine serum (FBS; 11560636, Gibco), 1% Glutamax (13462629, Gibco) and 1% antibiotics (11548876, Gibco), and after 24 hours the culture medium was replaced with osteogenic medium (Basal medium supplemented with 0,28 mM ascorbic acid (A4544, Sigma-Aldrich), 10 mM β-glycerol-2-phosphate (G9422, Sigma-Aldrich), and 10 nM dexamethasone (D4902, Sigma-Aldrich)). After 7 days of osteogenic induction an Alkaline Phosphatase Assay Kit (ab83371, Abcam) was used to estimate the osteogenic differentiation. Briefly, cells were detached from substrates with trypsin 0.05% (11590626, Gibco), counted and lysed. Next, lysates were centrifuged at 13.000 *g* for 3 minutes at 4ºC to remove insoluble materials. 4-methylumbelliferyl phosphate disodium salt (MUP) was added to the supernatants, which were then incubated for 30 minutes at 25ºC protected from light. Next, a stop solution was added and fluorescence intensity was measured in a plate reader with the excitation and emission wavelengths set at 360 and 440 nm, respectively. The resulting values were normalized against the cell count.

### Cell culture, immunostaining and image analysis

hBM-MSCs at an approximate density of 2.500-7.500 cells/cm^2^ depending on the specific experiment were seeded on the different surfaces in basal medium. Were specified, cultures were treated with 10 µM ROCK inhibitor Y-27632 dihydrochloride (Y0503, Sigma-Aldrich). At the given timepoints cells were fixed with 4% formaldehyde (28908, Sigma-Aldrich) for 15 minutes and permeabilized using 0.2% Triton X-100 (T9284, Sigma-Aldrich) at room temperature. Depending on the experiment, cells were stained with anti-vinculin (V9131, Sigma; 1:400), anti-ERK (ab184699, Abcam; 1:100) or anti-phospho-ERK (ab201015, Abcam; 1:100) antibodies for 1 hour at room temperature or with anti-YAP overnight at 4ºC (sc-101199, Santa Cruz Biotechnology; 1:300), followed by incubation with a fluorescently labelled secondary antibody (Alexa Fluor 488 anti-mouse IgG, A-11001, Molecular Probes, Life Technologies; 1:1000) for 1 hour at room temperature. Nuclei were counterstained with NucBlue (Hoechst 33342; R37605, Thermo Fisher) and the actin cytoskeleton with ActinRed 555 ReadyProbes (R37112, Thermo Fisher) following the manufacturer’s manual. Cells cultured on glass coverslips were used as control. Image acquisition was performed using a Nikon Ti-U inverted widefield microscope. For figure preparation, images were processed using Fiji (74) and for morphologic analysis images were processed using a custom-made pipeline of the CellProfiler software (Supplementary figure 1) (75).

### MYPT1 phosphorylation assay

After 24-hour culture, cells were trypsinized, counted, washed twice with cold PBS and collected by centrifugation at 2.500 *g* for 5 minutes. Pellets were resuspended in RIPA buffer (89900, Thermo Fisher) with protease inhibitors (11836170001, Sigma-Aldrich) and incubated for 15 minutes on ice. Samples were then centrifuged at 14.000 *g* for 15 minutes and the supernatant was collected. ROCK activity assay (CSA001, Sigma-Aldrich) was used following the manufacturer’s protocol. The results were expressed as total absorbance divided by the number of cells to estimate the average ROCK activity per cell in each experimental condition. Cells cultured on glass were used as reference conditions.

### RNA extraction and sequencing

After 24 hours of culture, total RNA was isolated using the RNAqueous-Micro Kit (AM1931, Thermo Fisher) following the manufacturer’s instructions. The quantity and quality of the RNAs were evaluated using Agilent RNA 6000 Pico Chips (5067-1513, Agilent Technologies). Sequencing libraries were prepared using “NEBNext Single Cell/Low Input RNA Library Prep Kit for Illumina” (E6420S, New England Biolabs) and “NEBNext Multiplex Oligos for Illumina (Index Primers 1-12)” (E7335S, New England Biolabs), following the instruction manual (Version 5.0_5/20). In brief, the protocol was started with 4 to 10 ng of total RNA and full-length cDNA was generated using a template switching method. Then, cDNA amplification and cleanup were performed and 1 μl of amplified cDNA was run on a DNA High Sensitivity Chip (5067-4626, Agilent Technologies) to assess cDNA quantity and quality. In the next step, fragmentation, end repair and A-tailing were carried out and adapters for Illumina were ligated to the amplified cDNA. Finally, library barcoding and amplification were achieved by PCR. Afterwards, libraries were quantified using Qubit dsDNA HS DNA Kit (Q32854, Thermo Fisher), visualized on an Agilent 2100 Bioanalyzer using Agilent High Sensitivity DNA kit (Agilent Technologies, 5067-4626) and sequenced in a NovaSeq 6000 (Illumina).

### Calcium imaging and signal quantification

After 24-hour adhering to the different β-PVDF surfaces, hBM-MSCs were stained with 2 μM Fluo4-AM (F14201, Thermo Fisher) for 30 minutes at 37ºC. Next, cultures were washed twice with PBS and stained with NucBlue (Hoechst 33342; R37605, Thermo Fisher) before adding DMEM/F12 HEPES without phenol red (11039-021, Gibco) for live imaging. Image sequences were acquired in a Nikon LV-IMA fluorescence microscope with the 10x objective, with appropriate filters for Fluo-4 and NucBlue, at 6-second intervals over a total of 16.5 minutes.

Next, we applied a shading correction and an intensity correction over time to each timelapse using the NIS-Elements software (Nikon) to allow the capture of local trends. In turn, the Linear Stack Alignment with SIFT Multichannel plugin (ImageJ) (76) was used to correct sample drift. Linear ROIs on the cell bodies were manually defined and their average intensity over the experiment duration was recorded. Next, the resulting intensity signal was linearly normalized and a moving average filter with a window size 4 was applied to facilitate comparisons across experiments. We set an intensity threshold of 75% above the baseline and maximal duration of 6 frames to identify transient increases in signal that represent biologically relevant events. To minimize false positives, we selected ROIs with at least three transient increases (peaks) and calculated their number and mean duration.

### Statistical analysis

Data is represented as average value +/- standard deviation (SD) of experimental replicas. To evaluate the statistical differences, data was analyzed assuming a Gaussian (normal) distribution. Except stated otherwise, equality of variances was not assumed across groups, therefore, differences between multiple groups were assessed using Welch’s ANOVA with the Brown–Forsythe test to account for unequal standard deviations. For post hoc multiple comparisons, the Dunnett’s T3 test was applied, using GraphPad Prism 6 (GraphPad Software, Inc.). p-values smaller than 0.1 are shown.

## Supporting information

Supplementary results and methods

## Data availability

RNA sequencing data have been submitted to NCBI’s Gene Expression Omnibus (GEO).

## Acknowledgements

The authors thank for technical and human support provided by SGIker (UPV/EHU/ ERDF, EU). This study was supported by Spanish Ministry of Science and Innovation (MCIN), project PID2022-138572OB-C42 with funding from MCIN/AEI/10.13039/501100011033 and FEDER, UE, and by the Health Department of the Basque Government [2023333036 and 2024333042]. RICORS: (RD21/00017/0024; RD24/0014/0025) Red Española de Terapias Avanzadas TERAV ISCIII funded by “Instituto de Salud Carlos III (ISCIII)” and co-funded by European Union (NextGenerationEU) “Plan de Recuperación Transformación y Resiliencia”. This study forms part of the Advanced Materials program and was supported by MCIN with funding from the European Union NextGenerationEU (PRTR-C17.I1) and by the Basque Government under the IKUR program. SMI was supported by a predoctoral scholarship provided by the Basque Government from the 2021 call.

## Credit author statement

SMI: Formal analysis, Investigation, Methodology, Visualization, Writing - original draft; YRV: Investigation; PRL: Investigation; LJR: Funding acquisition, Investigation, Methodology; CE: Funding acquisition and Investigation; NJR: Investigation; JA: Formal analysis, Investigation, Visualization; AMA: Investigation, Visualization; SLM: Conceptualization, Funding acquisition, Supervision, Writing - review and editing; US: Conceptualization, Formal analysis, Funding acquisition, Supervision, Writing - review and editing.

