## Supplementary results and methods for "Surface potential as a strong early osteogenic trigger via mechanotransduction and calcium accumulation"

**Supplementary information**

**Supplementary materials and methods**

**Western blotting**

Cell cultures were washed twice with cold PBS (10010-031, Gibco) and scraped in RIPA buffer (89900, Thermo Fisher) with protease (11836170001, Sigma-Aldrich) and phosphatase inhibitors (A32957, Thermo Fisher). Samples were collected, incubated for 15 minutes on ice and centrifuged at 12.000 *g* for 15 minutes. The supernatant was collected and the amount of protein present in the different samples was quantified using a BCA assay (71285-3, Sigma-Aldrich). Next, the volume equivalent to 5 μg of protein was pipetted into a test tube, and 5.8 μl of 6× loading buffer were added. The tubes were incubated at 100 °C for 5 minutes to denature the proteins. 12.5% polyacrylamide SDS-PAGE gels were run, and the protein wet transfer protocol was performed for 75 minutes at 400 mA, using a Criterion blotter (Bio-Rad). Membranes were blocked with 5% non-fat dry milk in Tris-buffered saline with Tween-20 (TBST) for 1 hour at room temperature and subsequently, incubated with primary antibodies (anti-phospho-ERK, ab201015, Abcam; 1:500; anti-ERK, ab184699, Abcam; 1:1000) in 1% non-fat dry milk in TBST overnight at 4 °C. Next, membranes were washes three times with TBST and incubated with the secondary antibody (Rabbit IgG; sc-525409, Santa Cruz Biotechnology; 1:2000) for 2 hours at room temperature. GAPDH ( sc-47724, Santa Cruz Biotechnology; 1:1000) was used as loading control. Finally, three washes with TBST were performed. To develop the membranes the surface of the blot was soaked in Immobilon Forte Western HRP substrate (WBLUF, Merck) for 1 minute. Afterwards, the signal was detected using an iBright CL1500 (MP Imaging System, Bio-Rad), and quantification was performed using the Fiji software.

**Supplementary figures**

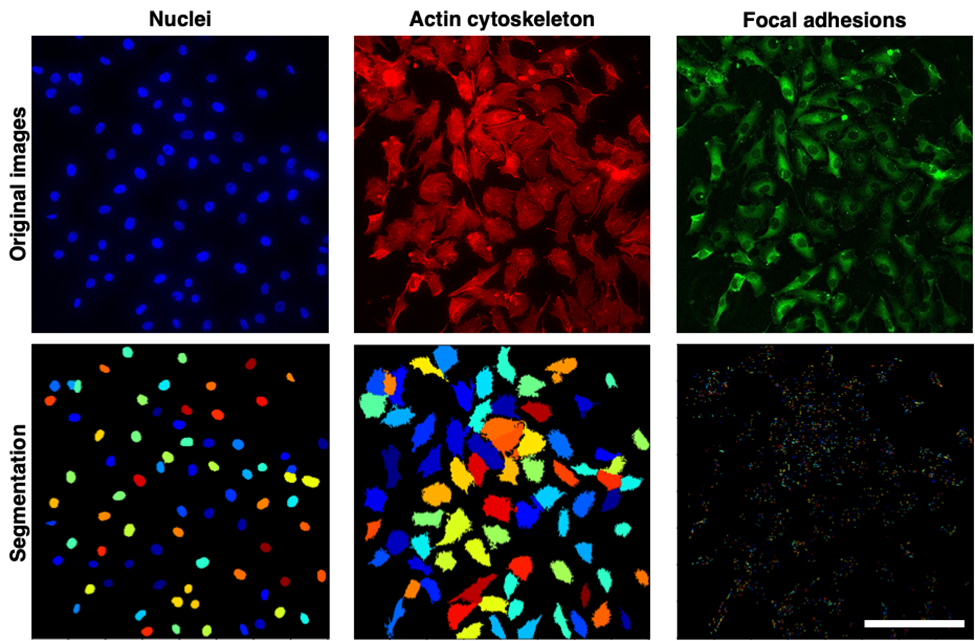

**Supplementary figure 1.- Image segmentation.** Fluorescence images of the hBM-MSCs labelled with Hoechst 33342, phalloidin and anti-vinculin (upper row) were segmented to determine nuclei size, spreading area and focal adhesion count (bottom row). Scale bar represents 200 µm.

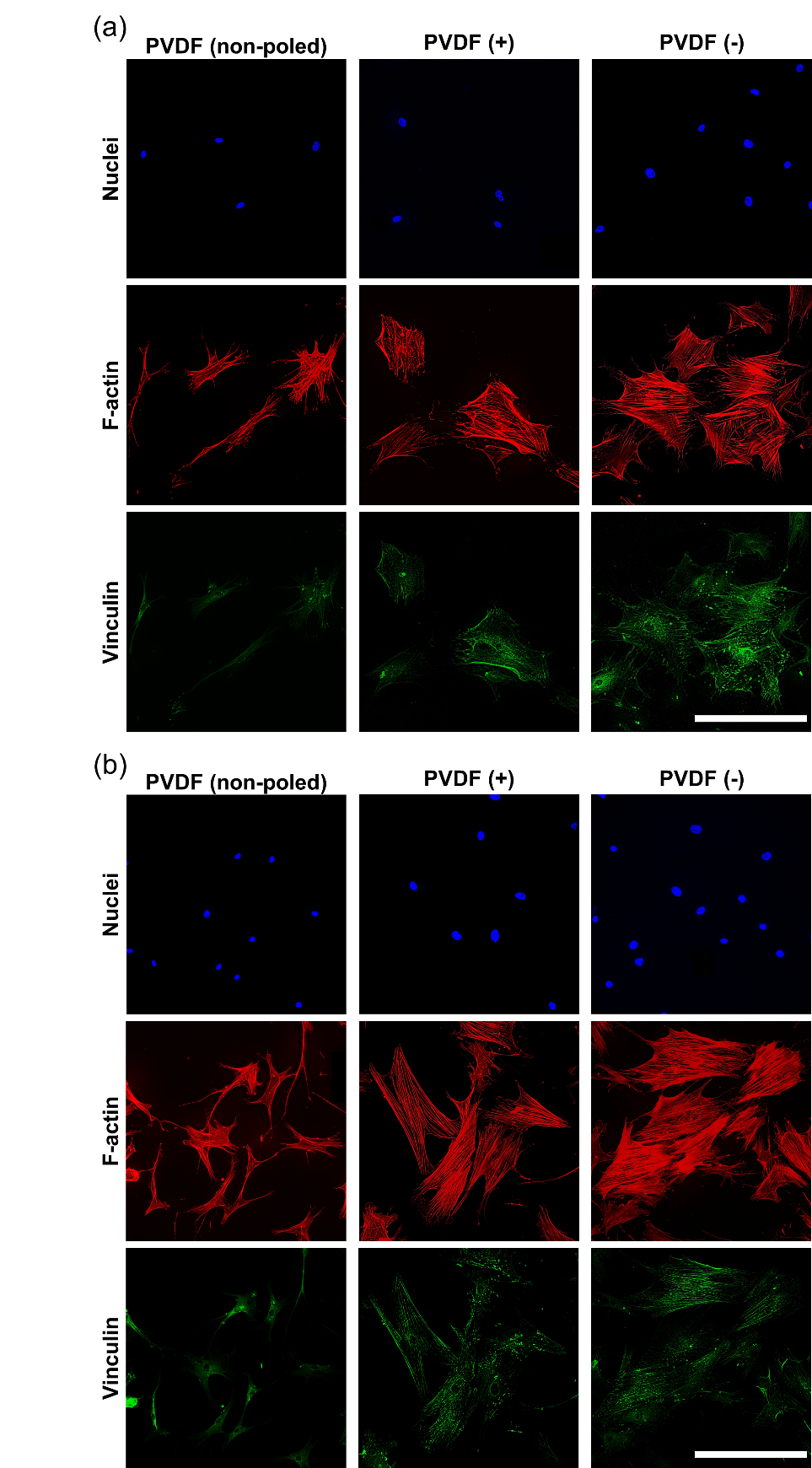

**Supplementary figure 2.- Fluorescence images of hBM-MSCs cultured on the different surfaces.**  Blue channel shows nuclei (stained with Hoechst 33342), green channel shows focal adhesions (stained with anti-vinculin), and the red channel shows F-actin (phalloidin) of hBM-MSc cultured for 24 hours (a) and 4 days (b) on the different surfaces coated with collagen type I. Scale bars represent 300 µm.

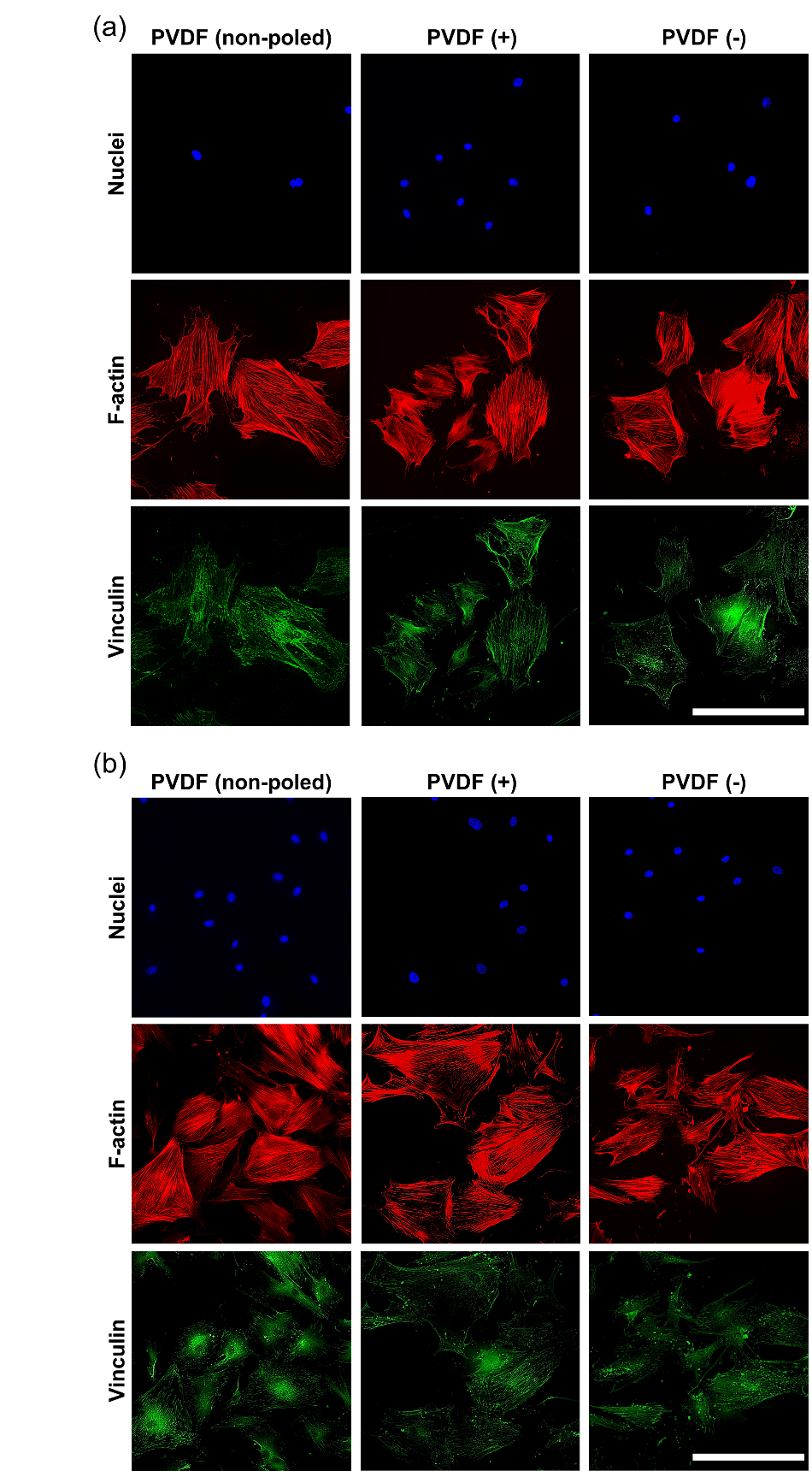

**Supplementary figure 3.- Fluorescence images of hBM-MSCs cultured on the different surfaces.**  Blue channel shows nuclei (stained with Hoechst 33342), green channel shows focal adhesions (stained with anti-vinculin), and the red channel shows F-actin (phalloidin) of hBM-MSc cultured for 24 hours (a) and 4 days (b) on the different surfaces coated with fibronectin. Scale bars represent 300 µm.

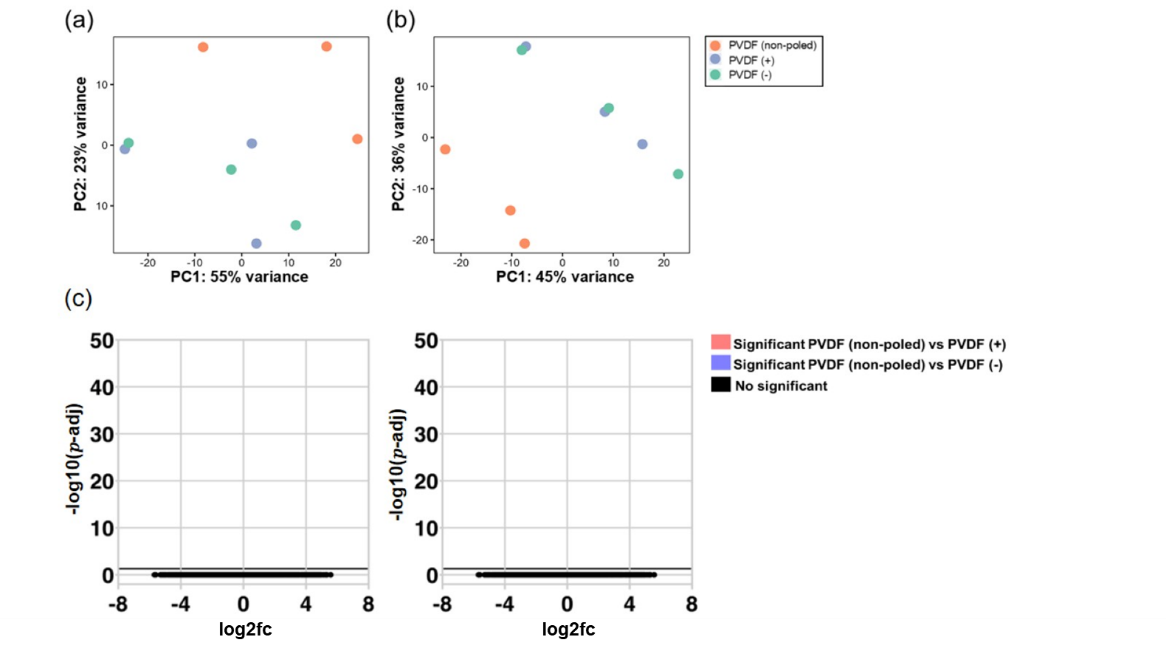

**Supplementary figure 4.- PCA analysis.** PC1 and PC2 variance of expression between the RNA-seq samples of cells adhering to the different β-PVDF films for 24 hours (a) and 4 days (b). (c) Volcano plots with the -log 10 (p-value) plotted against their respective log 2 (fold change) of genes differentially expressed in hBM-MSCs cultured for 24 hours (left panel) and 4 days (right panel) on β-PVDF (+) and β-PVDF (-) substrates.

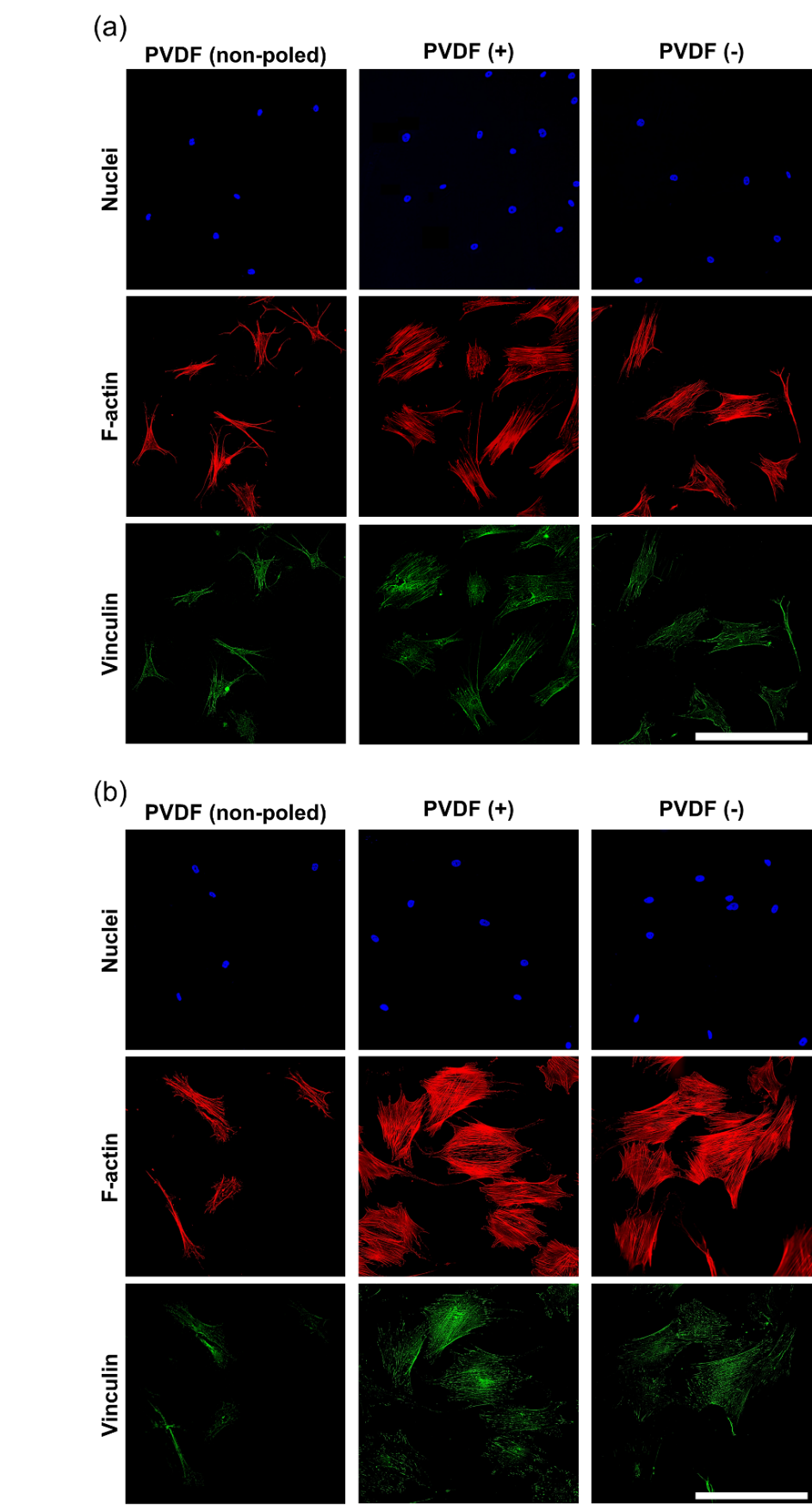

**Supplementary figure 5.- Fluorescence images of hBM-MSCs cultured on the different surfaces.**  Blue channel shows nuclei (stained with Hoechst 33342), green channel shows focal adhesions (stained with anti-vinculin), and the red channel shows F-actin (phalloidin) of hBM-MSc cultured for 24 hours (a) and 4 days (b) on the different surfaces coated with glutaraldehyde-fixed collagen type I. Scale bars represent 300 µm.

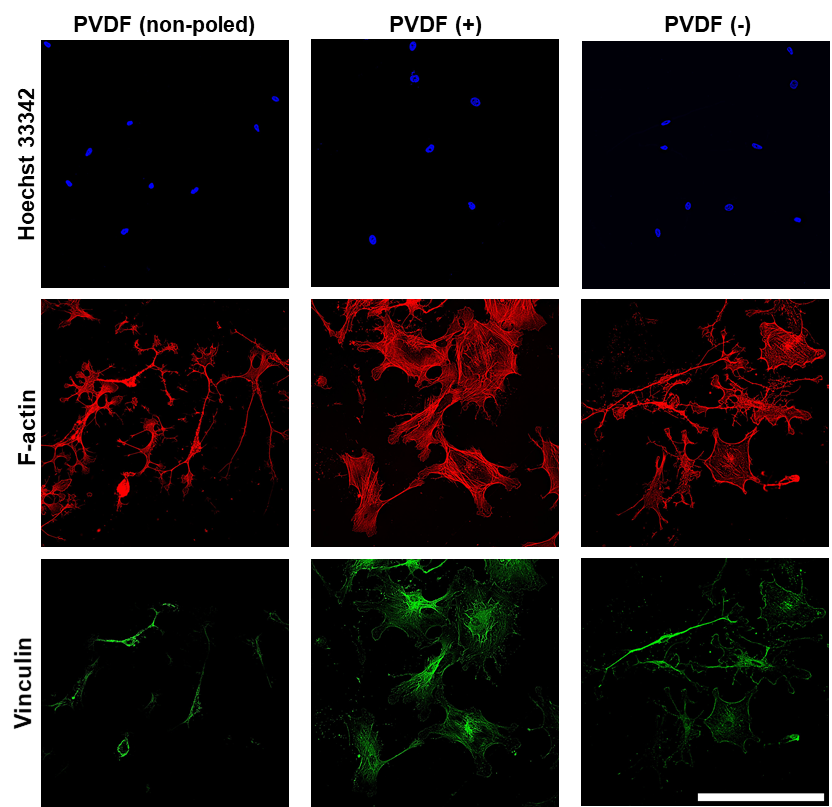

**Supplementary figure 6.- Fluorescence images of hBM-MSCs cultured on the different surfaces.**  Blue channel shows nuclei (stained with Hoechst 33342), green channel shows focal adhesions (stained with anti-vinculin), and the red channel shows F-actin (phalloidin) of Y-27632-treated hBM-MSc cultured for 24 hours on the different surfaces coated with collagen type I. Scale bar represents 300 µm.

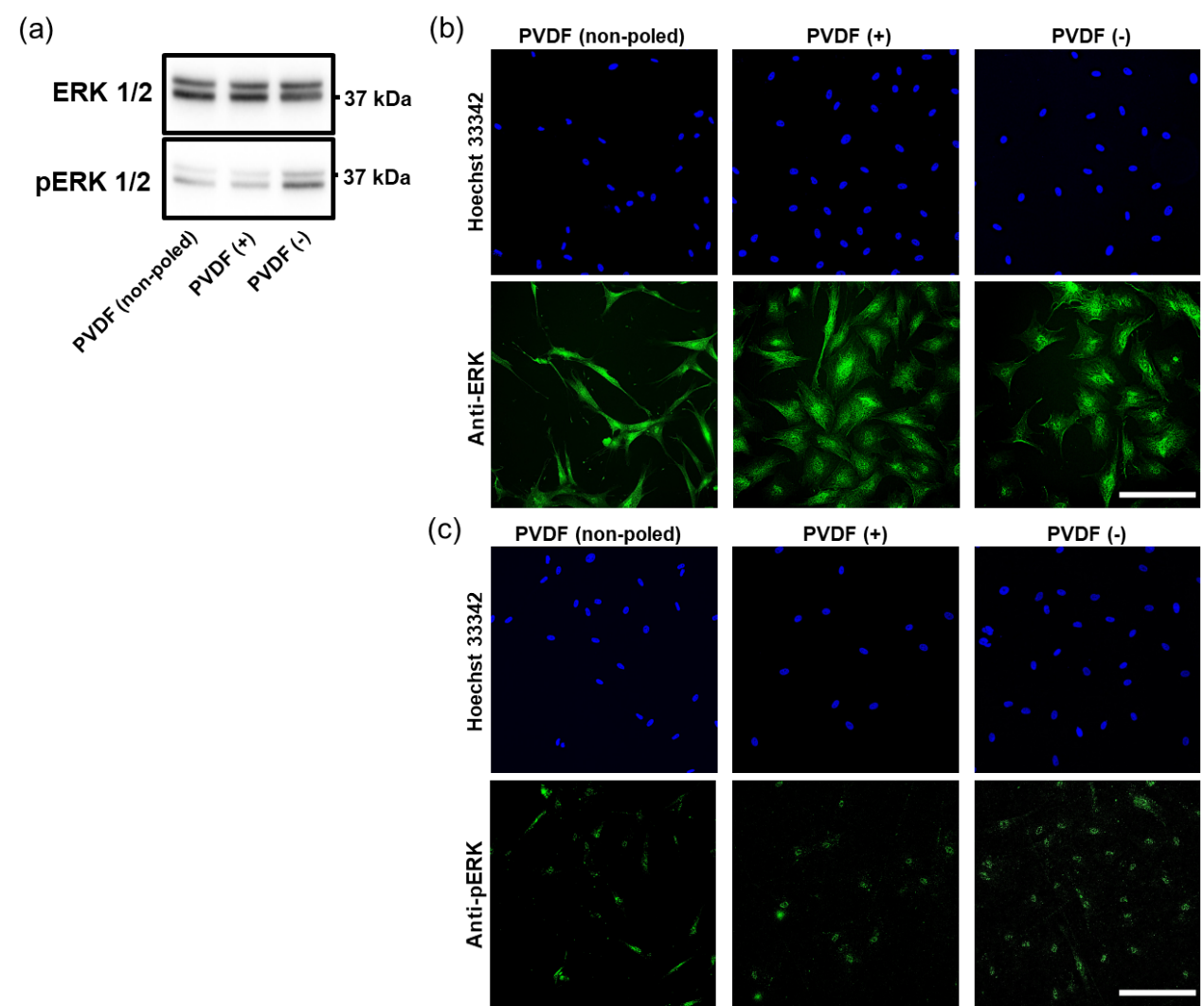

**Supplementary figure 7.- Analysis of ERK1/2 phosphorylation and subcellular localization.** (a) Western blot analysis of ERK1/2 phosphorylation. Total ERK (upper panel) and phosphorylated ERK (pERK, bottom panel). (b) Representative immunofluorescence images of hBM-MSCs stained for total ERK1/2 (in green) and (c) for pERK1/2. Nuclei were counterstained with Hoechst 33342. Scale bars represent 200 µm.
